# Temperature heterogeneity correlates with intraspecific variation in physiological flexibility in a small endotherm

**DOI:** 10.1101/2020.11.16.383877

**Authors:** Maria Stager, Nathan R. Senner, David L. Swanson, Matthew D. Carling, Douglas K. Eddy, Timothy J. Grieves, Zachary A. Cheviron

## Abstract

Phenotypic flexibility allows individuals to reversibly modify trait values and theory predicts an individual’s relative degree of flexibility positively correlates with the environmental heterogeneity it experiences. We tested this prediction by integrating surveys of population genetic and physiological variation with thermal acclimation experiments and indices of environmental heterogeneity in the Dark-eyed Junco (*Junco hyemalis*) and its congeners. We combined measures of thermogenic capacity for ~300 individuals, >21,000 single nucleotide polymorphisms genotyped in 192 individuals, and laboratory acclimations replicated on five populations. We found that *Junco* populations: (1) differ in their thermal performance responses to temperature variation *in situ*; (2) exhibit intra-specific variation in their thermogenic flexibility in the laboratory that correlates with heterogeneity in their native thermal environment; and (3) harbor genetic variation that also correlates with temperature heterogeneity. These results provide comprehensive support that phenotypic flexibility corresponds with environmental heterogeneity and highlight its importance for coping with environmental change.

## INTRODUCTION

Phenotypic plasticity — the ability of a single genotype to produce multiple trait values in response to an environmental cue — can be important for colonizing and persisting in novel environments^1–3^. As a result, the role of plasticity in adaptation to environmental variation has received significant attention^4–6^. These studies have documented standing genetic variation in plastic responses^7^, and the potential for plasticity to evolve in response to natural selection^8^. Because adaptive plasticity should increase fitness in variable environments, theory predicts that the magnitude of plasticity that individuals exhibit should positively correlate with the amount of environmental heterogeneity they experience^9–11^.

Empirical evaluations of this prediction have provided conflicting results, but most have focused on traits that are developmentally plastic (i.e., those that undergo environmentally-induced but irreversible changes to a trait value during specific developmental windows). For instance, morphological plasticity varies with diet breadth among ecotypes of threespine stickleback^12^. Similarly, plasticity in development time positively correlates with spatial variation in pool-drying regimes in the common frog (*Rana temporana*)^13^. Analogous patterns have been shown in plants as well. In a bindweed (*Convolvulus chilensis*), plasticity in leaf morphology and functional traits varies with interannual variation in rainfall^14^. Conversely, multiple studies have found that plasticity for thermal tolerance limits is not associated with latitudinal or thermal seasonality in *Drosophila*^15–17^. The disparities among these studies may arise from differences in the relationship between the temporal scale over which an environment varies and the time required for phenotypic induction. If so, traits that can be modified repeatedly may be expected to respond to intra-annual variation more strongly than traits that are irreversible.

Unlike developmental plasticity, phenotypic flexibility — the ability to reversibly modify trait values — provides repeated opportunities to match phenotypes to environmental change across an individual’s lifetime, especially in long-lived organisms^18^. Flexibility is predicted to evolve in environments characterized by frequent and predictable environmental variation^19^. Many morphological, physiological, and behavioral traits are flexible, and flexibility can represent an adaptive acclimatization response^20^. Determining the causes and consequences of variation in flexibility among individuals is therefore crucial to our understanding of adaptive evolution, and our ability to predict species’ resilience to environmental change. Yet few empirical studies have tested whether geographic variation in the degree of flexibility is associated with spatial patterns of environmental heterogeneity (but see ref.^21^), and none have accounted for non-independence among populations (due to shared common ancestry and ongoing gene flow), which can obscure true relationships among variables of interest when analyzing intraspecific patterns of phenotypic variation^22^.

If environmental heterogeneity influences variation in adaptive plasticity/flexibility across a species’ range, we would therefore also expect to find that environmental heterogeneity structures population genetic variation as well. Patterns of environmentally segregating genetic variation can arise when local selective regimes influence the rates of gene flow among environmentally distinct habitats^23,24^ and are not uncommon among widespread species^25,26^. No study, however, has tested whether the same environmental driver correlates with both genetic variation and variation in the degree of flexibility in a trait across a species’ entire range.

To uncover the environmental drivers of geographic variation in phenotypic flexibility, we investigated the flexible capacity of a key physiological trait in a temperate songbird, the Dark-eyed Junco (*Junco hyemalis*). Juncos are particularly well suited to investigations of phenotypic flexibility due to the extensive phenotypic variation they exhibit and the broad range of environments they occupy^27^. The *J. hyemalis* lineage is comprised of five distinct morphotypes and 14 subspecies that inhabit a variety of habitats (Fig. S1) and exhibit conspicuous differences in life history, migratory tendency, physiology, size, song, plumage, and behavior^27–29^. This diversity is thought to have arisen since the most recent glacial maxima when the *J. hyemalis* ancestor diverged from *J. phaeonotus fulvescens* of southern Mexico and subsequently expanded its range across North America^30,31^. While environmental factors have been shown to partition genetic variation within a subset of *J. hyemalis* taxa^32^, the major *J. hyemalis* morphotypes do not exhibit strong population genetic differentiation^31^ suggesting that considerable phenotypic diversity persists in the face of high rates of gene flow. The role that environmental conditions play in driving the diversification of this lineage thus remain unclear.

Juncos have also been the subject of intense physiological study. Many *J. hyemalis* groups winter at temperate latitudes, and temperate environments place a premium on endogenous heat production in small homeothermic endotherms to maintain a relatively constant body temperature^33^. As a result, lab and field studies show that juncos increase their thermogenic capacity (the ability to generate heat; quantified as the peak metabolic rate under cold exposure, M_sum_) in the cold^34–36^. This heightened thermogenic capacity is associated with a reduced risk of hypothermia for juncos^36,37^, with a failure to achieve adequate thermogenic output having dire consequences for organismal fitness in small endotherms^38–40^. Increases in thermogenic performance are also accompanied by changes occurring at lower hierarchical levels of biological organization^35,41–43^. Thus, a higher thermogenic capacity may be energetically costly to maintain due to the additional metabolic machinery required to achieve elevated thermogenic performance^44^. Phenotypic flexibility in thermogenic capacity could therefore help mediate a balance between thermoregulation and its associated maintenance costs in response to fluctuating selective pressures^45^. Nonetheless, it is unknown whether an individual’s capacity for thermogenic flexibility is influenced by the degree of thermal variability it experiences throughout the year.

We drew upon natural variation that exists across the *Junco* distribution to understand geographic variation in flexibility of thermogenic performance, and to test for associations between environmental heterogeneity, phenotypic flexibility, and population genetic structure. To do this, we first surveyed *in situ* geographic variation in *Junco* thermogenic capacity to determine which environmental indices structure variation in this trait. We then characterized fine-scale, range-wide population genetic structure within the *Junco* genus to determine whether it is influenced by the same climatic indices. Finally, we performed a laboratory acclimation experiment on five *Junco* populations that differ in their annual thermal regimes to test whether environmental heterogeneity predicts the degree of thermogenic flexibility. This approach allowed us to perform the first study that combines measures of physiological flexibility with indices of climatic variation while controlling for nonindependence among populations. We predicted that junco populations that experience greater seasonal temperature variation would exhibit higher thermogenic flexibility than those from more thermally stable regions. By combining these approaches, our results shed light on the ecological conditions that promote the evolution of increased flexibility and address long-standing hypotheses in the field of evolutionary biology.

## RESULTS AND DISCUSSION

### Geographic variation in thermogenic performance

Although several studies have characterized broad-scale *inter*specific patterns in endothermic thermogenic performance^46–48^, far less is known about the potential for or the underlying environmental correlates of *intra*specific variation in thermogenic performance (but see refs. ^49–51^). We assayed M_sum_ for 292 juncos at 8 sites across the U.S. and correlated recent weather conditions with patterns of *in situ* variation (Fig. 1). The number of individuals, number of sampling days, seasons, and years varied across sites with 86 total site-days of environmental variation and 5 morphotypes included in our dataset (Table S1).

**Figure 1.**
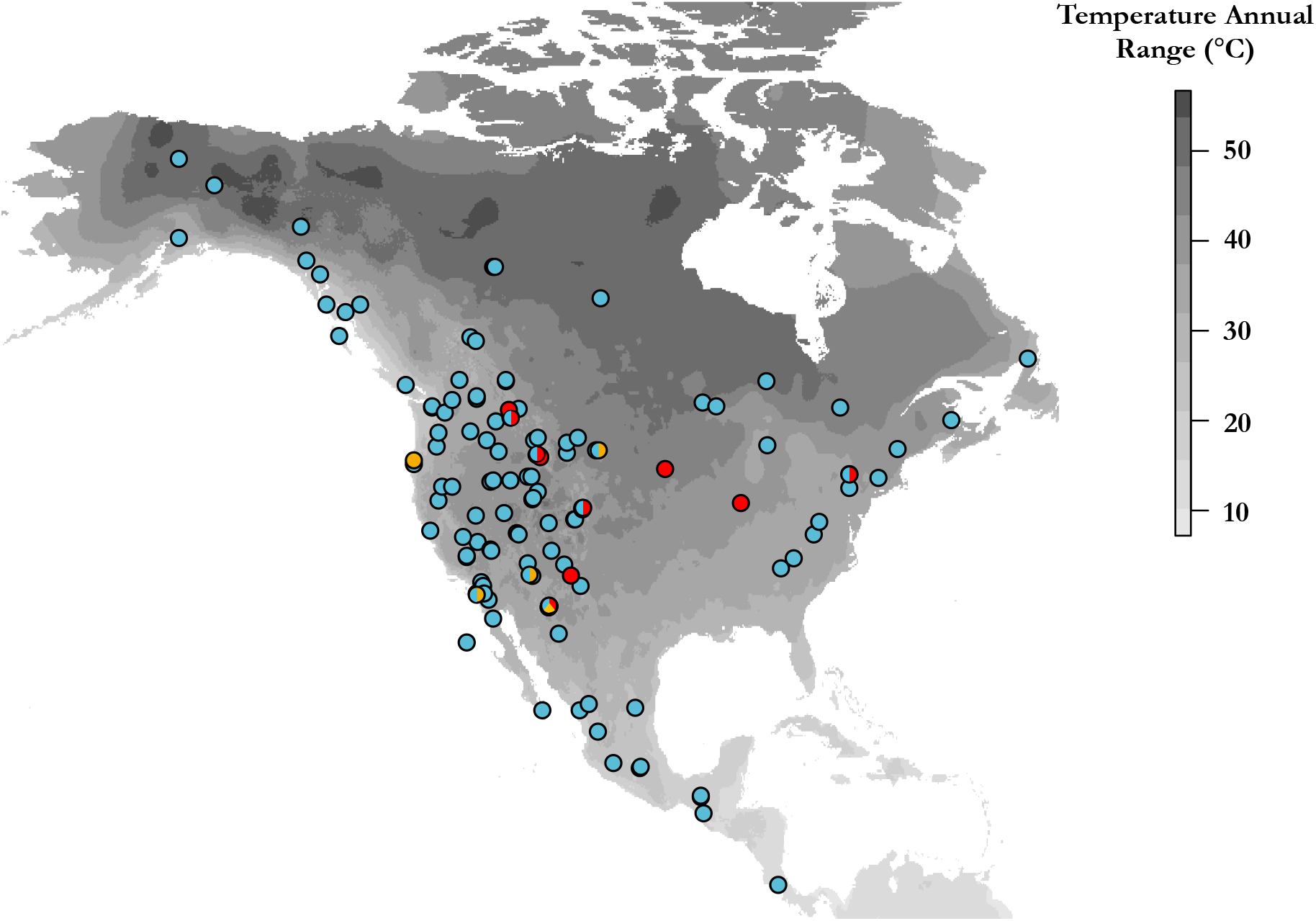
Temperature annual range of North America and Central America from the WorldClim dataset (grayscale) with junco sampling locations for *in situ* measurements (red), phylogeography (blue), and acclimations (yellow).

Geographic variation in M_sum_ corresponded to environmental variation (Table S2). The most well-supported model included body mass (Mb), daily temperature range (T_d_range_), morphotype, and a T_d_range_× morphotype interaction and explained 50% of the variation in M_sum_ (AICc = 883.4, *w_i_* = 0.998). While Mb positively correlated with M_sum_ (*β* = 1.39 ± 0.17, *p* < 0.001), the model also showed a persistent effect of morphotype on M_sum_ after controlling for differences in Mb, with Oregon Juncos exhibiting the lowest and Slate-colored Juncos the highest M_sum_ values (Table S3). This could suggest local adaptation in thermogenic performance among populations. However, juncos that experienced larger T_d_range_ in the 7-8 d prior to capture also had the highest M_sum_, indicating that they may be responding to short-term heterogeneity in their thermal environment (Table S3). This is consistent with recent laboratory findings showing that *J. h. montanus* can make substantial changes to M_sum_ within one week of exposure to low temperature36. Moreover, we also found that the inclusion of the T_d_range_× morph interaction term substantially improved model fit indicating that some populations respond differentially to temperature variation in the wild (Fig. 2). Taken together, this suggests that *Junco* populations may differ in their physiological flexibility and that variation in the temperature range across their distribution may play an important role in shaping this flexibility.

**Figure 2.**
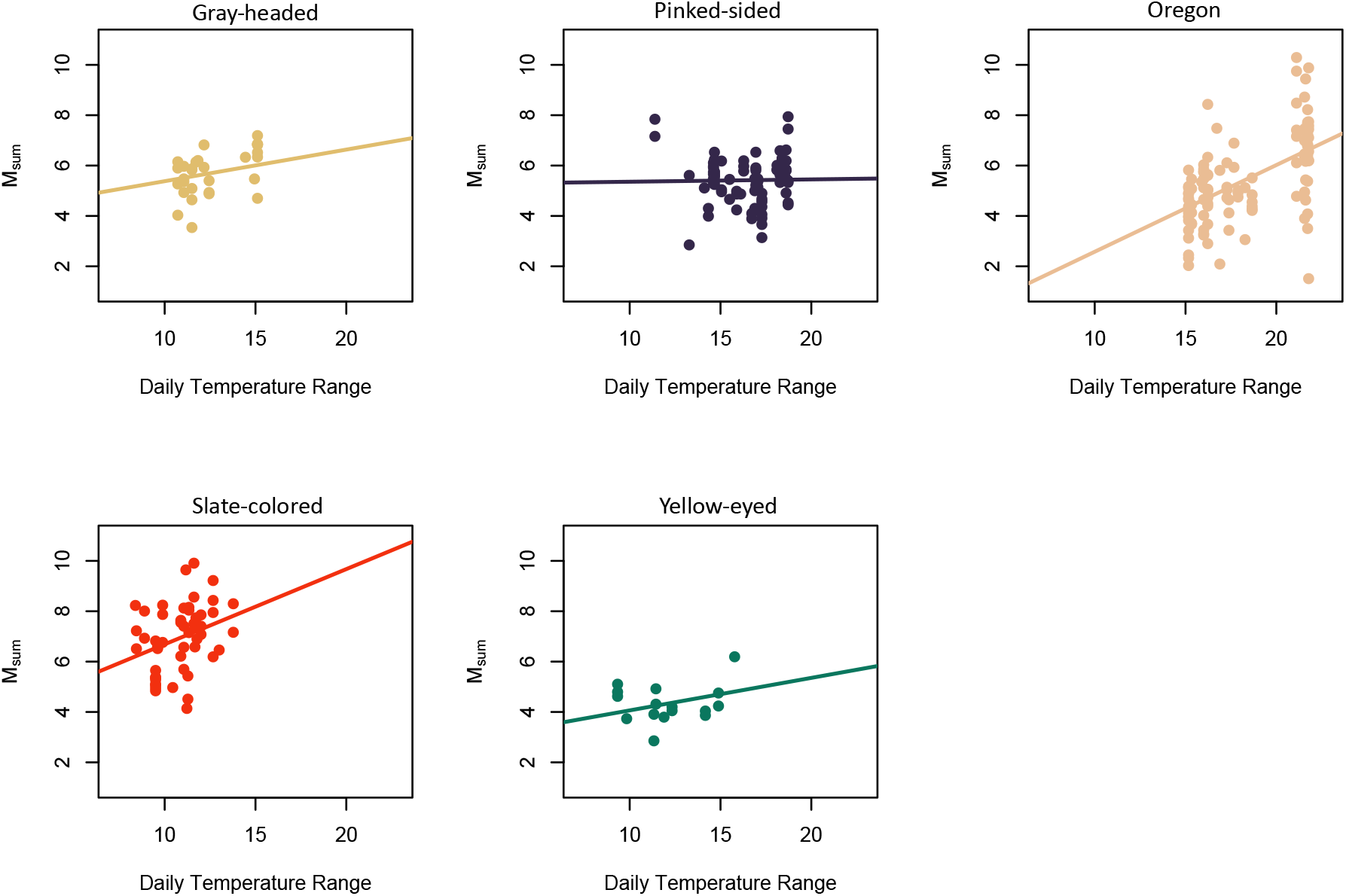
Effect of daily temperature range on *in situ* M_sum_ while controlling for differences in M_b_. Model results shown in Table S3.

### Genotype-environment associations

The structure of the environment can also influence rates of gene flow among habitats and is therefore an important component determining the selective regime acting on local populations^23,24^. To understand how environmental variables might correspond to *Junco* population genetic variation, we genotyped 21,971 biallelic SNPs from 181 individuals that were selected to maximize geographic and environmental variance while representing all recognized *Junco* species/subspecies (Fig. S1). We first used a principal components analysis (PCA) to assess levels of underlying genetic variation across the radiation and found that the first two PC axes explained 7.1% of the total genetic variance. Major clusters identified by the PCA corresponded to the ‘Sky Island’ lineages of central America (*J. vulcani*), southern Mexico (*J. p. alticola* and *J. p. fulvescens*), and southern Baja (*J. p. baird*i), as well as the Guadalupe Junco (*J. insularis*), with all other morphotypes grouping together (Fig. 3). The fact that all *J. hyemalis* and northern *J. phaeonotus* taxa comprised one overlapping genetic cluster likely reflects the rapid expansion of this lineage over the last ~20,000 years^30,31^.

**Figure 3.**
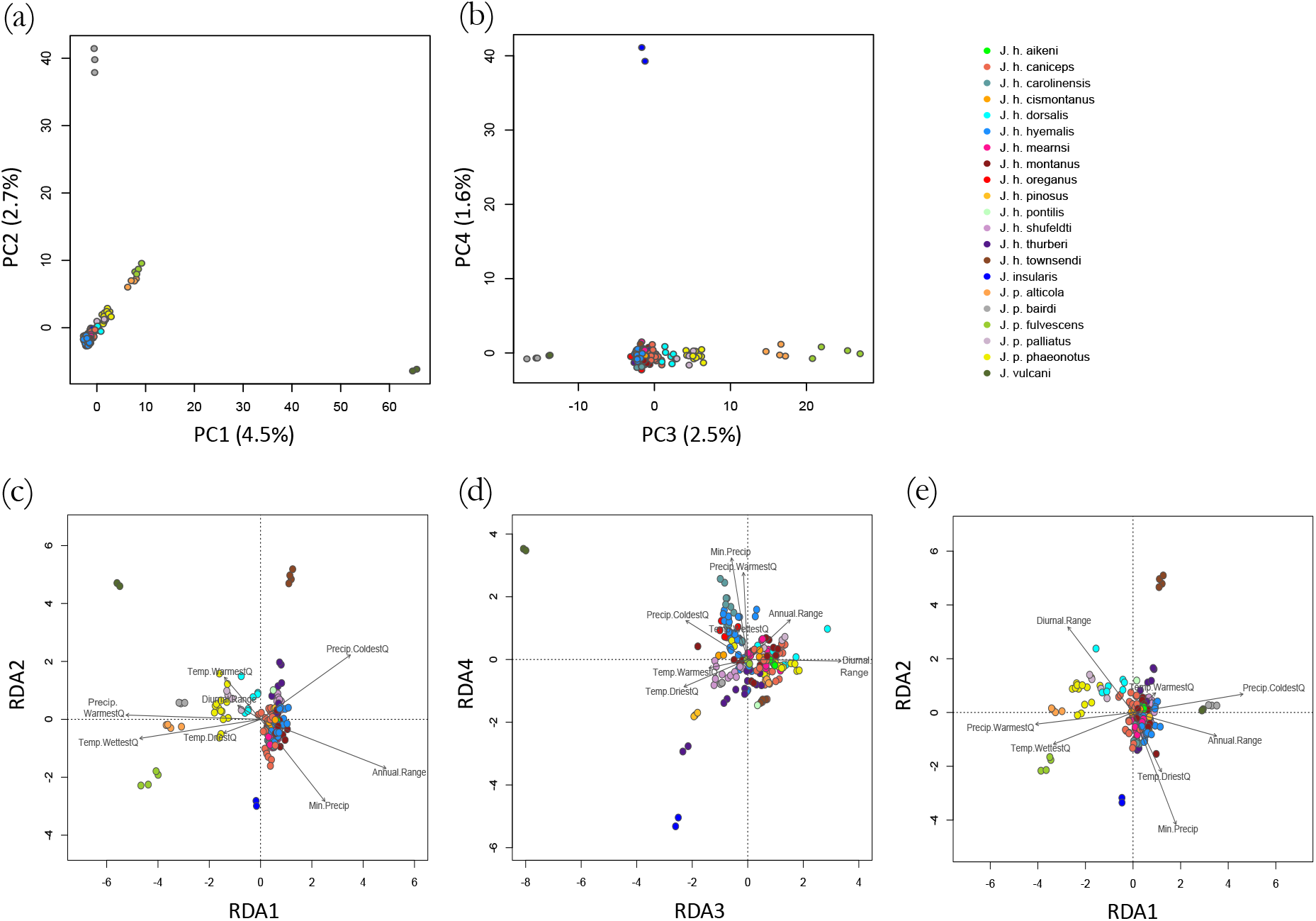
Population genetic structuring along 4 PC axes (a,b), 4 RDA axes in unconditioned RDA (c,d), and 2 RDA axes in a conditioned model (e). Arrows indicate loadings of 8 WorldClim variables. Dots represent individuals, colors denote museum-based taxon assignments.

In instances where populations are not clearly distinguishable and environmental gradients are continuous, genotype-environment association methods can aid in the detection of signatures of natural selection^52^. In particular, redundancy analysis (RDA) is a powerful multivariate tool for identifying even weak correlations between genetic and environmental data^53^. We thus performed an RDA to quantify the population genetic variance that partitions with climatic indices while controlling for background genetic structure. Eight RDA axes explained 6.2% of the total genetic variance across *Junco* in our unconditioned model and 5.6% in our conditioned model controlling for background population structure. Permutation tests confirmed the significance of the model in both cases (p = 0.001). The first 4 RDA axes were significant in the unconditioned model (p < 0.05), whereas the first 3 RDA axes were significant in the conditioned model. A variance partition analysis showed that temperature annual range explained more genetic variation than any other climatic variable that we tested in both the unconditioned (0.52%) and conditioned models (0.60%; Table 1). Additionally, we detected 751 outlier SNPs exhibiting associations with the first 3 axes in the conditioned model. Of these, 128 and 48 outlier SNPs corresponded most strongly to mean diurnal temperature range and annual temperature range, respectively. This complements our above finding that *in situ* physiological variation is also influenced by temperature range. However, it is not known whether these sites are involved in conferring flexibility and additional studies are necessary to reveal the genomic architecture underlying thermogenic flexibility.

**Table 1.** Genetic variance across 21,971 SNPs partitioned by 8 climatic variables in an unconditioned RDA and an RDA conditioned on background structure.

| **Variable** | **Unconditioned**<br>**Adj. R<sup>2</sup>** | **Conditioned**<br>**Adj. R<sup>2</sup>** |
| --- | --- | --- |
| Diurnal Temperature Range | 0.0026 | 0.0034 |
| Temperature Annual Range | 0.0052 | 0.0060 |
| Temperature Wettest Quarter | 0.0012 | 0.0012 |
| Temperature Driest Quarter | 0.0022 | 0.0015 |
| Temperature Warmest Quarter | 0.0008 | 0.0008 |
| Minimum Precipitation | 0.0028 | 0.0029 |
| Precipitation Warmest Quarter | 0.0037 | 0.0030 |
| Precipitation Coldest Quarter | 0.0034 | 0.0035 |

Our results also show that several taxa overlap in orthogonal space (Fig. 3). Recent work by Friis et al.^32^ instead found that Oregon Junco taxa exhibited distinct environmental partitioning across their distribution. One reason for this difference may be that we included more *Junco* taxa in this study. In addition, Friis and colleagues sampled each taxon from a single site, which tends to confound environmental and geographic distance^54^. Our dataset encompasses far more environmental variation across varying geographic distances, and thus uniquely highlights the role of seasonal and diurnal climatic variation in structuring population genetic variation within and among *Junco* taxa.

### Flexible responses to temperature acclimation treatments

To formally test whether phenotypic flexibility in thermogenic capacity correlated with environmental heterogeneity, we performed an acclimation experiment using individuals from five populations across the western U.S. Our ability to connect *Junco* populations to native climatic regimes is restricted by our limited knowledge of junco movements across the year, and we therefore focused on populations that likely remain resident to one narrow geographic area in order to reliably reconstruct their climatic histories. Based on the results of our *in situ* sampling, we selected focal populations to maximize variation in the annual temperature range (T_range_) they experienced. Genetic differentiation among populations (F_ST_) ranged from 0.019 to 0.051 (Table S4), while pairwise environmental and genetic distances did not covary (partial Mantel test: *r* = −0.40, *p* = 0.82) allowing us to simultaneously tease apart the effects of both factors.

Prior to acclimation, temperature treatment groups did not differ in M_sum_ (*t* = −1.10, df = 89, *p* = 0.27) or M_b_ (*t* = 0.04, df = 89, *p* = 0.97). However, both traits did positively correlate with native temperature range (M_sum_: *β* = 0.07 ± 0.01, *p* < 0.001; M_b_: *β* = 0.25 ± 0.02, *p* < 0.001). We therefore controlled for individual differences in pre-acclimation M_sum_ and M_b_ in our subsequent analysis. Over the course of the experiment we found that ΔM_sum_ (measured as the difference between post- and pre-acclimation measures) was higher in larger and cold-acclimated birds (M_b_: posterior mean = 0.53, 95% CI = [0.03, 1.00], *p* = 0.03; Cold Treatment: posterior mean = 0.62, 95% CI = [0.35, 0.88], *p* < 0.001). We also found an interaction between native T_range_ and treatment, such that populations from more variable climates exhibited the greatest increase in M_sum_ in the cold (posterior mean = 0.60, 95% CI = [0.07, 1.14], *p* = 0.03), while populations from less variable climates exhibited little or no change in M_sum_ (Fig. 4). Our results therefore clearly support our prediction that juncos from more thermally variable environments exhibit a greater capacity for thermogenic flexibility.

**Figure 4.**
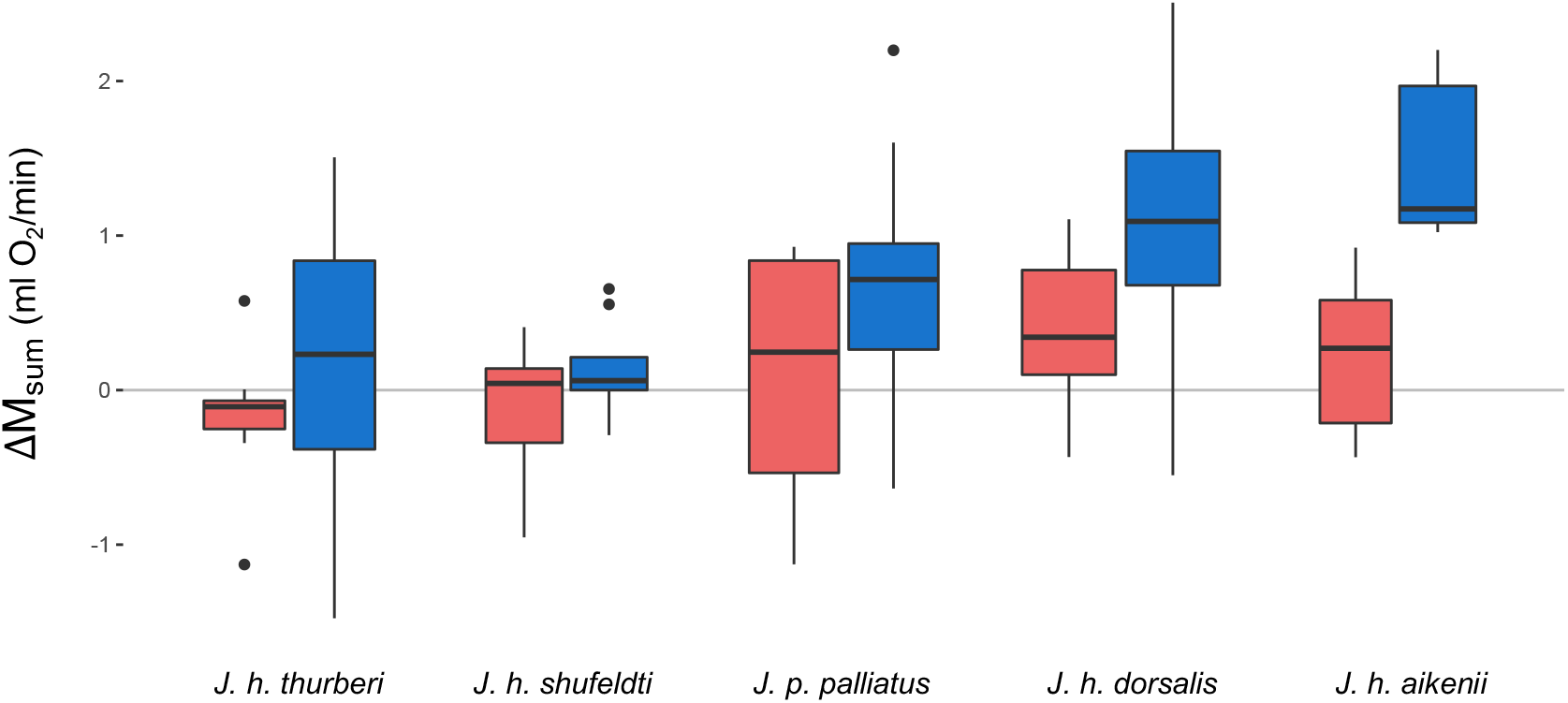
Change in M_sum_ (post- minus pre-acclimation) over a three-week acclimation period for each population ordered from lowest to highest native temperature range (from left to right). Control birds in red, cold-acclimated birds in blue; *n* = 95.

## Conclusions

Our multifaceted approach integrated measures of population genetic variation with whole-organism measures of physiological performance in both the lab and field, as well as indices of environmental variation to uncover the environmental drivers of geographic variation in phenotypic flexibility. We provide evidence that temperature variation drives patterns of intra-specific variation in thermal performance and found that junco populations responded differentially to weather cues *in situ*. This pattern was also replicated in the laboratory where thermogenic flexibility in juncos correlates with the heterogeneity of their native thermal environment. Moreover, range-wide population genetic variation also correlates with ambient thermal variation, providing evidence that environmental heterogeneity may be an important selective force driving junco population divergence. Together, these results suggest that physiological flexibility may play a key role in local adaptation in this broadly distributed genus.

While this result supports theory^9–11^, it contrasts with previous empirical studies in ectotherms that found that plasticity/flexibility in thermal physiology did not correspond to environmental heterogeneity^15–17,55^. While the cause of this disparity is unclear, one aspect that has largely been overlooked in previous studies is the role of historical demographic processes in shaping adaptive plasticity/flexibility. Gene flow^56^, colonization history^57^, population size^58^, and the standing genetic variation of founding individuals^59^ are all important factors shaping adaptive outcomes^26^. For example, although gene flow can constrain adaptive divergence^56^, high gene flow among selective regimes is predicted to favor increased plasticity/flexibility in order to aid offspring that experience a dissimilar environment from their parents^3,10,60^. Comprehensive studies that simultaneously incorporate both contemporary ecological conditions and population demographic processes are necessary to fully understand the role of environment heterogeneity in structuring plasticity/flexibility.

There are also several biological differences between ectotherms and endotherms that may contribute to disparities in evolutionary patterns of phenotypic plasticity/flexibility. In general, many ectotherms rely on behavioral thermoregulatory mechanisms and possess a number of avoidance strategies (e.g., diapause, hibernation, or migration) that may be used to buffer against environmental extremes^61^. Thus, the amount of thermal heterogeneity that an individual or population experiences may not correspond to broad-scale climatic patterns across the year. Endotherms maintain a relatively constant body temperature in comparison, despite sometimes large temperature differentials with their ambient environment^33^. While many endotherms also exhibit hibernation and migratory behaviors, small songbirds that reside in temperate regions year-round, like juncos, are particularly exposed to thermal heterogeneity^45^. These differences may lead to divergent selection pressures on flexibility and thermal performance traits among taxonomic groups.

This study greatly expands our knowledge of endothermic responses to environmental variation and their capacity for thermal acclimatization. Understanding flexibility in organismal thermal tolerances is important for predicting population dynamics^62^, making habitat delineations^63^, and modeling disease transmission^64^, all of which are especially relevant in light of ongoing global climate change. Although many recent macrophysiological approaches characterizing potential organismal responses to climatic change employ a single metric of thermal tolerance, effectively treating tolerance as a trait that is canalized and invariant across a species’ range (e.g., refs ^55,65,66^), our results highlight the capacity for populations to vary geographically in their physiological response to environmental cues. When coupled with datasets like ours, biophysical models that incorporate intraspecific patterns in acclimatization will improve our ability to predict organismal responses to climate warming.

## METHODS

### *In situ* data and analysis

#### In situ sampling

We captured juncos by mist net or potter trap at sites in Arizona (breeding season), Colorado (breeding), Illinois (wintering), Montana (breeding), New Mexico (wintering), New York (breeding), South Dakota (wintering), and Wyoming (breeding), spanning 16° in latitude and 37° in longitude (Fig. 1; Table S1). We classified individuals into known morphotypes based on plumage (Gray-headed, Oregon, Pink-sided, Slate-colored, and Yellow-eyed). These morphs have distinct breeding distributions that often encompass the ranges of multiple subspecies^27,28,67^; however, the wintering ranges overlap for many morphs.

#### In situ metabolic assays

We assayed M_sum_ of captured birds using open-flow respirometry near the site of capture. All measurements were made within 48 h of capture to avoid the effects of captivity on metabolic rates, though most were completed within 24 h. M_b_ was quantified before each measurement began. M_sum_ trials were conducted during the birds’ light phase. A single individual was placed in a metabolic chamber in a dark, temperature-controlled environment. We pumped dry, cooled heliox gas (21% O_2_, 79% He) through the animal’s chamber at a constant flow rate. We subsampled the outflow current, dried it (Drierite), scrubbed it of CO_2_ using ascarite, and dried it again before quantifying the O_2_ concentration using a FoxBox (Sable Systems). Trials were conducted using static cold exposure (−5°C) for CO, IL, MT, NM, NY, and WY birds and sliding cold exposure (starting at 0°C) for AZ and SD birds; however, both methods have been shown to produce similar estimates of M_sum_^68^. Trials lasted until O_2_ consumption plateaued or declined for several minutes. We also sampled a blank chamber before and after trials to account for potential fluctuations in baseline, ambient air.

We used custom R scripts to quantify M_sum_ as the highest instantO_2_ consumption averaged over a 5-min period (https://github.com/Mstager/batch_processing_Expedata_files). We discarded measures characterized by large drift in baseline O_2_ (owing to ambient temperature fluctuations affecting the Fox Box) or inconsistent flow rates resulting in a total sample size of *n* = 292 individuals. Following measurements, birds were subject to different fates: either released, exposed to acclimation experiments^36^, or immediately euthanized and deposited in museums (Table S1). Metabolic data from SD have been previously published^69^.

#### Environmental data for in situ sampling sites

To account for an individual’s recent acclimatization history, we retrieved weather data associated with each collection site (rounded to the nearest hundredth of a degree latitude/longitude) from the DayMet dataset using the R package *daymetr*^70^. This dataset is composed of daily weather parameter estimates derived using interpolation and extrapolation from meteorological observations for 1km x 1km gridded surfaces across North America^71^. We downloaded daily estimates of minimum temperature (T_min_), maximum temperature (T_max_), precipitation (prcp), water vapor pressure (wvp), and daylength (dayl) for the 14 d prior to and including each individual’s capture date. We additionally calculated daily temperature range (T_d_range_) as T_max_ – T_min_. We selected a conservative potential acclimatization window because, given their migratory nature, we do not know how long juncos were present at a site before sampling occurred, preventing us from incorporating extensive windows (e.g., months). Prior work in *J. hyemalis* suggested a window of 7-14 d would be appropriate^36,72^. We therefore calculated running averages for each weather variable across acclimatization windows varying from 7-14 d preceding capture. We also retrieved elevation (elev) for each site using the package *googleway*^73^.

#### Analyses for in situ data

All analyses were conducted in R version 4.0.2^74^. To determine whether junco M_sum_ varied with environmental variation, we constructed seven linear models with M_b_, morphotype (morph), and a single environmental variable (elev or running averages of T_min_, T_max_, T_d_range_, prcp, dayl, or wvp) as main effects. We first standardized continuous predictor variables according to ref.^75^ using the package *arm*. We then used AICc values to evaluate differences in model fits among environmental variables and with that of a null model (including only M_b_ and morph as predictors) in order to identify a single ‘best’ environmental variable and acclimatization window explaining variation in M_sum_. As an indicator of differences in flexibility, we additionally tested for an interaction between the best environmental variable and morph to determine if populations differed in their response to environmental cues. To quantify the effect sizes of the interaction term, we reran the model with each morph as the reference.

### Population genetic data

#### Sampling, sequencing, and SNP generation

For phylogeographic reconstruction, we obtained muscle tissue samples (*n* = 192) from museum specimens collected across the breeding distribution of all *Junco* species and subspecies (Table S5). This included 2-30 individuals per taxonomic unit sampled from 94 geographic localities representing the majority of U.S. counties, Canadian provinces, and Mexican states for which tissue samples exist (Fig. S1). We then employed restriction-site-associated DNA (RAD)-sequencing, which offers a reduced representation of the genome that can be mined for thousands of single-nucleotide polymorphisms (SNPs) among individuals^76,77^.

We extracted whole genomic DNA from each sample using a Qiagen DNeasy Blood and Tissue extraction Kit and prepared RAD-libraries according to ref.^78^ (protocol ver. 2.3 downloaded from dryad: https://datadryad.org/stash/dataset/doi:10.5061/dryad.m2271pf1). Briefly, we digested whole genomic DNA with two restriction enzymes (*EcoRI* and *Mse1*), ligated adaptor sequences with unique barcodes (8-10 bp) for each individual, performed PCR amplification, and then performed automated size selection of 300–400 bp fragments (Sage Science Blue Pippen). We split paired geographic samples between the two libraries such that all taxa were represented in each library of 96, pooled individuals. Libraries were sequenced on separate flow-cell lanes of an Illumina HiSeq 4000 at UC Berkeley’s V.C. Genomics Sequencing Lab, yielding over 300 million, 100-nt single-end reads per lane.

We demultiplexed reads, retained reads with intact *EcoRI* cut sites, removed inline barcodes and adapters, and performed quality filtering (removed Phred score < 10) using *process_radtags* in STACKS ver. 2.1^79^. This resulted in final reads of *μ* = 92 bp in length and a mean of 2.05 million reads per individual. We removed 3 individuals in each lane that failed to sequence (< 100,000 reads/individual, comprising 5 *J. hyemalis* and 1 *J. insularis*). We used *bwa mem*^80^ to align reads to the *J. h. carolinensis* genome^32^, which we downloaded from NCBI (Accession GCA_003829775.1). An average of 91% of reads mapped and mapping success did not differ among *Junco* species.

We executed the STACKS pipeline to call SNPs and exported one SNP per locus for loci present in all four *Junco* species in vcf format *(populations: −p* 4 *--write-random-snp*), resulting in 178,574 SNPs. We further filtered the dataset using vcftools (ver. 0.1.16, Danecek et al. 2011). We removed sites with mean depth of coverage across all individuals < 5 (*--min-meanDP*) and > 50 (*--max-meanDP*), minimum minor allele count < 3 *(--mac*), more than 50% missing data (--*max-missing*), and characterized by indels *(--remove-indels*). We then removed 5 *J. hyemalis* individuals with > 60% missing data (*--remove*). Finally, we removed sites with > 5% missing data (--*max-missing*) and filtered sites that departed from Hardy-Weinburg equilibrium assessed by species (*p* < 0.001), resulting in 21,971 biallelic SNPs across 181 individuals. We exported the dataset in plink.raw format for downstream analysis.

#### Genotype-environment association analyses

To determine if environmental variation corresponded to genetic variation across *Junco*, we employed the SNP dataset in a redundancy analysis (RDA). RDA is a multivariate ordination technique that has been used to identify multiple candidate loci and several environmental predictors simultaneously^53,82^. Because RDA requires no missing data, we first imputed data for *n* = 68,311 missing sites (1.7% of total sites) using the most common genotype for the species at each site. We downloaded 19 interpolated monthly climate variables corresponding to the geographic origin of each specimen from WorldClim^83^ at a resolution of 2.5’ using the R package *raster* ver. 3.3-13^84^. We centered and standardized climate variables according to ref.^75^. We excluded highly correlated variables (*r* ≥ 0.70) resulting in 8 variables retained: mean diurnal temperature range (BIO_2_, a measure of monthly temperature variation), temperature annual range (BIO7), mean temperature of the wettest quarter (BIO8), mean temperature of the driest quarter (BIO9), mean temperature of the warmest quarter (BIO10), precipitation of the driest month (BIO14), precipitation of the warmest quarter (BIO18), and precipitation of the coldest quarter (BIO19). We used these 8 climatic variables as predictors in RDAs executed with the package *vegan* ver. 2.5-6^85^. We performed both a simple RDA with no conditional treatment and a partial RDA conditioned on background population structure. For the latter, we summarized population genetic structure using the first two principal components from a principal component analysis of the imputed SNP data with the R package *ade4* ver 1.7-15^86^. In both RDAs, we tested for, but did not find, multicollinearity among predictor variables (variance inflation factor < 5 for all variables). We assessed the significance of the RDA models and of the constrained axes using an ANOVA-like permutation technique in *vegan* at *p* ≤ 0.05 (*n* = 999 and *n* = 99 permutations, respectively). We estimated the proportion of variance explained by each climatic variable using variance partitioning^87^. We then identified candidate SNPs for environmental adaptation as those outside of a 3 standard deviation cutoff from the mean loading and characterized each candidate SNP by the predictor variable with which it had the strongest correlation (per ref.^53^).

### Acclimation experiments

#### Population sampling for acclimation experiments

We combined information gained from a literature search, eBird sightings, and expert opinion (*pers. comm.* Tom Martin) to identify five focal populations for phenotypic sampling that (1) were likely to be non-migratory, (2) represented different morphological subspecies, and (3) maximized variation in annual temperature range within the United States. These populations include the White-winged Junco (*J. h. aikeni*) of the Black Hills, a coastal population of Oregon Junco (*J. h. shufeldti*), a highland population of Red-backed Junco (*J. h. dorsalis*), a sky island population of Yellow-eyed Junco (*J. p. palliatus*), and a well-studied, urban population of Oregon Junco (*J. h. thurberi*)^88^. However, it is possible that some of these populations exhibit seasonal, altitudinal migrations within their geographic area of residence, e.g., *J. p. palliatus*^89^.

We captured ≤ 25 individuals from each focal population. Capture periods differed for each population in order to increase the likelihood that targeted individuals were resident year-round, as well as due to time and permitting constraints. For instance, one partially migratory population (*J. h. aikeni*) with distinct morphological features was caught in the winter to increase the likelihood that the individuals used were non-migratory. The other four populations, which bred in areas where other, morphologically similar juncos over-winter, were captured in the breeding season when other subspecies were not present. Specifically, *J. h. shufeldti* (*n* = 20) were captured 14-15 July 2018 in Coos and Douglas Counties, OR; *J. p. palliatus* (*n* = 24) were captured 27 July 2018 in Cochise County, AZ; *J. h. dorsalis* (*n* = 25) were captured 30-31 July 2018 in Coconino County, AZ; *J. h. aikeni* (*n* = 15) were captured 6-9 March 2019 in Lawrence County, SD; and *J. h. thurberi* (*n* = 20) were captured 22-26 July 2019 in San Diego County, CA. In spite of these differences, we do not think capture season likely influenced acclimation ability because: (1) we subjected all individuals to a long adjustment period in order to erase prior acclimatization history before the experiment began (see below); and (2) the addition of a season term in our best model explaining variation in M_sum_ *in situ* did not improve model fit, suggesting that juncos are responding to proximate cues rather than making seasonal adjustments (*β*_season_ = −0.11, *p* = 0.64, ΔAIC_c_ = 1.97).

#### Acclimation treatments

Within days of capture, we ground-transported all birds to facilities at the University of Montana where birds were housed individually under common conditions (23°C with 12 h dark: 12 h light) for ≥ 8 weeks (*μ* = 63 d, range = 57-71 d). We had previously determined that a period of 6 wk is sufficient to reduce variation in metabolic traits among individuals (Fig. S2). Following this adjustment period, we assayed M_sum_ (see below). We allowed birds ~24 h to recover and then randomly assigned individuals from each population into treatment groups and exposed them to either cold (3°C) or control (23°C) temperatures. Treatments lasted 21 d in duration. Constant 12 h dark: 12 h light days were maintained for the duration of the experiment, and food and water were supplied *ad libitum*. The diet consisted of a 2:1 ratio by weight of white millet and black oil sunflower seed, supplemented with ground dog food, live mealworms, and water containing vitamin drops (Wild Harvest D13123 Multi Drops). These experimental conditions were chosen based on previous work in *J. h. hyemalis* exposed to the same temperatures, which revealed substantial increases in M_sum_ over the same duration^35^.

Brood patches and cloacal protuberances were not present after the adjustment period. At the end of treatments, we euthanized individuals using cervical dislocation. Gonads, identified during dissection, were regressed in all but one individual (a male *J. h. dorsalis*) post-acclimation.

Eight individuals died during the capture-transport and adjustment periods (1 *J. h. dorsalis,* 4 *J. h. shufeldti*, 1 *J. h. thurberi,* and 2 *J. p. palliatus*). Additionally, one *J. h. thurberi* individual exhibited lethargy upon introduction to the cold treatment, died within the first 24 h of cold acclimation, and was removed from analyses. This resulted in a total sample size of *n* = 95 individuals (Table S6).

#### Metabolic assays for acclimation experiments

We quantified M_sum_ in a temperature-controlled cabinet using open-flow respirometry both before and after acclimation treatments as described above. We measured M_b_ immediately before each assay. M_sum_ trials were conducted at −5°C for pre-acclimation measures and −15°C for post-acclimation measures. Although this experimental temperature difference could have elicited higher post-acclimation M_sum_, we did not find variation in M_sum_ between the two time points in Control birds, suggesting that the two procedures provided similar levels of cold challenge (paired t-test: *p* = 0.60). Because trials occurred at various times throughout the day, we tested for, but did not find, a linear effect of trial start time on M_sum_ either before or after acclimation (*p_pre_* = 0.54, *p_post_* = 0.94).

#### Climate data for acclimation populations

We reconstructed the annual thermal regime experienced by a population using interpolated monthly climate data downloaded from the WorldClim dataset^83^ with package *raster*. Specifically, we extracted the temperature annual range variable (BIO7) at a resolution of 2.5’ for the site of capture (hereafter T_range_). Focal populations varied in T_range_ by 21°C (Fig. 1).

#### Pair-wise genetic distance of acclimated populations

To quantify genetic differentiation among acclimated populations, we extracted whole genomic DNA from the muscle tissue of acclimated individuals and prepared RAD-libraries as above with an additional PCR step (per protocol v. 2.6)^78^. We pooled all 95 individuals into a single flow-cell lane of an Illumina HiSeq X for sequencing at Novogene Corporation Inc. yielding ~1 billion, 150-nt paired-end reads. We demultiplexed reads, removed inline barcodes and adaptors, and performed quality filtering (removed Phred score < 10) using *process_radtags*. This resulted in final reads of *μ* = 147 bp in length and a mean of 7 million reads per individual. We removed 6 individuals that sequenced poorly (< 200,000 reads/individual, comprising 1 *J. h. dorsalis,* 1 *J. p. palliatus,* 1 *J. h. shufeldti*, and 3 *J. h. thurberi*). We aligned reads to the *J. h. carolinensis* genome with *bwa mem* (51% of reads mapped). We then sorted, merged, and removed read duplicates with samtools ver. 1.3^90^. We used the STACKS pipeline to call SNPs from loci present in all 5 populations and exported one SNP per locus in vcf format resulting in 282,929 SNPs (*populations*: *-p 5 −-write-random-snp*).

We filtered the dataset and estimated patterns of genetic differentiation using *vcftools*. We removed sites with mean depth of coverage across all individuals < 20 (*--min-meanDP*) and > 300 (*-- max-meanDP*), minor allele frequency < 5% (*--min-maf*), and > 10% missing data (--*max-missing*), and removed indels (*--remove-indels*), resulting in 1069 biallelic SNPs across 89 individuals. Finally, we estimated pairwise F_ST_ among the focal taxa by calculating weighted Weir’s theta^91^.

#### Analyses for acclimation data

We first performed a partial mantel test to ascertain that environmental and genetic distances do not covary among our sampling sites. We calculated pairwise genetic distance as F_ST_/(1-F_ST_). We estimated pairwise environmental differences among the five sites as the Euclidean distance for 5 WorldClim variables (after removing redundant variables at *r* ≥ 0.70 from the original 19 WorldClim variables). We simultaneously controlled for geographic distance, estimated as pairwise geodesic distance among sampling sites with the package *geosphere*^92^. We then employed these matrices of pairwise genetic, environmental, and geographic distance in a partial mantel test with the package *vegan*^85^.

We verified that phenotypic differences did not exist among treatment groups before acclimations began using t-tests for M_b_ and M_sum_. We also used linear regression to quantify the effects of temperature range on pre-acclimation trait values. To evaluate whether environmental variation corresponded with flexibility, we related climatic data to M_sum_ for each population while simultaneously incorporating population demography. We used Markov Chain Monte Carlo generalized linear mixed models that allow for Bayesian approaches^22,93^ with the *MCMCglmm* package^94^. We constructed a model to explain variation in thermogenic flexibility (ΔM_sum_; post-minus pre-acclimation M_sum_) with pre-acclimation M_b_, temperature treatment, T_range_, and treatment × T_range_ interaction as main effects and pairwise F_ST_ as a random effect. We standardized all continuous predictor variables according to^75^. We used default priors and ran the model for 1,000,000 iterations with a burn-in of 10,000 and a thinning interval of 100. We examined the resulting trace plots to verify proper convergence.

## ETHICS

This work was completed with approval from the U.S. Fish and Wildlife Service (MB84376B-1 to M.S.; MB01543B-0 to Z.A.C.; MB758442 to D.L.S.; MB06336A-4 to M.D.C.; MB45239B-0 to T.J.G.; MB757670-1 to David Winkler; and MB094297-0 to Christopher Witt,), the State of Arizona Game and Fish Department (E19253811 to M.S. and SP590760 and SP707897 to D.L.S.), the State of California Department of Fish and Wildlife (13971 to M.S.), Colorado Parks and Wildlife (10TRb2030A15 to M.D.C.), the Illinois Department of Natural Resources (NH13.5667 to Z.A.C.), the Montana Department of Fish Wildlife and Parks (2016-013 and 2017-067-W to M.S.), the New Mexico Department of Game & Fish (#3217 to Christopher Witt), the New York State Division of Fish, Wildlife, & Marine Resources (LCP 1477 to David Winkler), the Oregon Department of Fish and Wildlife (108-18 to M.S.), the State of South Dakota Department of Game, Fish, and Parks (19-13 to M.S. and 06-03, 07-02, 08-03 to D.L.S.), Wyoming Game and Fish (754 to M.D.C.), and the Institutional Animal Care and Use Committees at Cornell University (2001-0051 to David Winkler), the University of Illinois (13385 to Z.A.C.), the University of Montana (010-16ZCDBS-020916 and 030-18ZCDBS-052918 to Z.A.C.), the University of South Dakota (03-08-06-08B to D.L.S), and the University of Wyoming (A-3216-01 to M.D.C.).

## ACKNOWLEDGMENTS

We are indebted to the many natural history museums that contributed tissue loans to this project, including John Klicka and the University of Washington Burke Museum, the American Museum of Natural History, the Cleveland Museum of Natural History, the Cornell University Museum of Vertebrates, the Field Museum of Natural History, the Louisiana State University Museum of Natural Science, the Midwest Museum of Natural History, the Museum of Southwest Biology, the Museum of Vertebrate Zoology, the New York State Museum, the Royal Alberta Museum, the San Diego Natural History Museum, the Smithsonian National Museum of Natural History, the University of Alaska Museum, the University of Montana Zoological Museum, and the University of Wyoming Museum of Vertebrates. We are very thankful to the many hands that provided help and logistical assistance collecting birds: Gregory Toreev, Cole Wolf, Trey Sasser, Phred Benham, Nick Sly, Henry Pollock, Phil Unitt, Chris Witt, Andy Johnson, Blair Wolf, Eric Gulson, David Winkler, Charles Dardia, Link Smith, Pamela Yeh, Eleanor Diamant, Kevin Burns, and Point Loma Nazarene University. We are grateful to the field stations that hosted our work: UNM Sevilleta Field Station, UW-NPS Research Station, Mt. Evans Field Station, and the Southwestern Research Station. We also thank Brett Addis, Rena Schweizer, and Thom Nelson for guidance with RAD processing and pop gen analysis, and the Cheviron lab for feedback on an earlier version of this manuscript.

## AUTHOR CONTRIBUTIONS

M.S. and Z.A.C. conceived of the study; M.D.C., D.K.E., and N.R.S. helped perform field work; D.L.S. performed *in situ* measurements in AZ and SD; T.J.G. provided sampling permits; M.S. performed all other data collection and analyses and drafted the manuscript; all authors contributed edits to the manuscript.

## DATA AVAILABILITY

Raw reads have been deposited in the NCBI Sequence Read Archive (PRJNA678344). Phenotypic data are included in the Supplementary Materials. All associated scripts are deposited on github (https://github.com/Mstager/junco_flexibility_scripts).

