## Supplemental Tables and Figures for "Temperature heterogeneity correlates with intraspecific variation in physiological flexibility in a small endotherm"

**Supplemental Materials**

**Table S1**. *In situ* individuals. Includes individual ID, morphotype, capture locality (state, latitude, longitude), capture date, measurer, mass, M_sum_, and voucher catalog number (where appropriate). Provided as .csv file.

**Table S2.** Effects of environmental variables on junco M_sum_ *in situ*. Weather variables are mean value for the indicated number of days preceding capture. All acclimatization windows are reported for T_d_range_, while only the best window is reported for other variables for brevity. All models include M_b_ and morph as covariates and all continuous variables were standardized; *n* = 292 individuals.

| **Variable** | **Window (d)** | **K** | **Deviance** | **AIC_c_** | $\boldsymbol{\Delta}$**AIC_c_** |
| --- | --- | --- | --- | --- | --- |
| T_d_range_ | 0-8 | 3 | 347.50 | 895.98 | 0 |
| T_d_range_ | 0-7 | 3 | 348.90 | 897.15 | 1.17 |
| T_d_range_ | 0-10 | 3 | 351.93 | 899.68 | 3.70 |
| T_d_range_ | 0-9 | 3 | 352.11 | 899.83 | 3.85 |
| T_d_range_ | 0-11 | 3 | 354.80 | 902.05 | 6.07 |
| T_d_range_ | 0-12 | 3 | 358.95 | 905.45 | 9.47 |
| prcp | 0-7 | 3 | 359.00 | 905.48 | 9.5 |
| T_d_range_ | 0-13 | 3 | 360.86 | 906.99 | 11.01 |
| T_d_range_ | 0-14 | 3 | 362.48 | 908.30 | 12.32 |
| T_max_ | 0-7 | 3 | 405.59 | 941.12 | 45.14 |
| T_min_ | 0-7 | 3 | 425.65 | 955.22 | 59.24 |
| wvp | 0-12 | 3 | 426.65 | 955.90 | 59.92 |
| elev |  | 3 | 427.48 | 956.46 | 60.48 |
| null |  | 2 | 430.72 | 956.56 | 60.58 |
| dayl | 0-14 | 3 | 429.84 | 958.07 | 62.09 |

**Table S3.** Effects of T_d_range_, morph, and their interaction on *in situ* M_sum_ while controlling for differences in M_b_. T_d_range_ is the running average for the 8 d preceding and including the capture date. All continuous variables were standardized. Estimates vary depending on which morphotype is used as the reference. AICc = 883.38, *n* = 292 individuals, df = 281, R^2^ = 0.50.

*PS as reference:	GH as reference:*

| **Variable** | **Beta** | **SD** | **p** |
| --- | --- | --- | --- |
| **Intercept** | **5.52** | **0.14** | **< 2.0 x 10^-16^** |
| **M_b_** | **1.39** | **0.17** | **2.79 x 10^-14^** |
| **Morph (SC)** | **1.98** | **0.57** | **5.87 x 10^-4^** |
| **Morph (YE)** | **-1.41** | **0.54** | **9.70 x 10^-3^** |
| Morph (GH) | 0.61 | 0.42 | 0.15 |
| **Morph (OR)** | **-0.65** | **0.20** | **1.55 x 10^-3^** |
| T_d_range_ | 0.07 | 0.49 | 0.89 |
| **T_d_range_ x SC** | **2.10** | **0.94** | **0.03** |
| T_d_range_ x YE | 0.87 | 1.07 | 0.41 |
| T_d_range_ x GH | 0.84 | 0.99 | 0.39 |
| **T_d_range_ x OR** | **2.43** | **0.56** | **2.11 x 10^-5^** |

| **Variable** | **Beta** | **SD** | **p** |
| --- | --- | --- | --- |
| **Intercept** | **6.13** | **0.40** | **< 2.0 x 10^-16^** |
| **M_b_** | **1.39** | **0.17** | **2.79 x 10^-14^** |
| Morph (PS) | -0.61 | 0.42 | 0.15 |
| **Morph (SC)** | **1.37** | **0.67** | **0.04** |
| **Morph (YE)** | **-2.01** | **0.65** | **2.23 x 10^-4^** |
| **Morph (OR)** | **-1.25** | **0.43** | **4.00 x 10^-3^** |
| T_d_range_ | 0.91 | 0.85 | 0.29 |
| T_d_range_ x PS | -0.84 | 0.99 | 0.39 |
| T_d_range_ x SC | 1.26 | 1.17 | 0.28 |
| T_d_range_ x YE | 0.03 | 1.28 | 0.99 |
| T_d_range_ x OR | 1.59 | 0.89 | 0.07 |

*YE as reference:	SC as reference:*

| **Variable** | **Beta** | **SD** | **p** |
| --- | --- | --- | --- |
| **Intercept** | **4.11** | **0.52** | **6.91 x 10^-14^** |
| **M_b_** | **1.39** | **0.17** | **2.79 x 10^-14^** |
| **Morph (GH)** | **2.04** | **0.65** | **2.23 x 10^-3^** |
| Morph (OR) | 0.76 | 0.55 | 0.17 |
| **Morph (PS)** | **1.41** | **0.54** | **9.70 x 10^-3^** |
| **Morph (SC)** | **3.38** | **0.75** | **9.05 x 10^-6^** |
| T_d_range_ | 0.94 | 0.95 | 0.32 |
| T_d_range_ x GH | -0.03 | 1.28 | 0.98 |
| T_d_range_ x OR | 1.56 | 0.99 | 0.11 |
| T_d_range_ x PS | -0.87 | 1.07 | 0.41 |
| T_d_range_ x SC | 1.23 | 1.24 | 0.33 |

| **Variable** | **Beta** | **SD** | **p** |
| --- | --- | --- | --- |
| **Intercept** | **7.49** | **0.55** | **< 2.0 x 10^-16^** |
| **M_b_** | **1.39** | **0.17** | **2.79 x 10^-14^** |
| **Morph (GH)** | **-1.37** | **0.67** | **0.04** |
| **Morph (OR)** | **-2.62** | **0.58** | **9.94 x 10^-6^** |
| **Morph (PS)** | **-1.98** | **0.57** | **5.87 x 10^-4^** |
| **Morph (YE)** | **-3.38** | **0.74** | **9.05 x 10^-6^** |
| **T_d_range_** | **2.17** | **0.80** | **7.31 x 10^-3^** |
| T_d_range_ x GH | -1.26 | 1.17 | 0.28 |
| T_d_range_ x OR | 0.33 | 0.85 | 0.70 |
| **T_d_range_ x PS** | **-2.10** | **0.94** | **0.02** |
| T_d_range_ x YE | -1.23 | 1.24 | 0.33 |

*OR as reference:*

| **Variable** | $\boldsymbol{\beta}$ | **SD** | ***p*** |
| --- | --- | --- | --- |
| **Intercept** | **4.87** | **0.15** | **< 2.0 x 10^-16^** |
| **M_b_** | **1.39** | **0.17** | **2.79 x 10^-14^** |
| **Morph (GH)** | **1.25** | **0.43** | **4.00 x 10^-3^** |
| **Morph (PS)** | **0.65** | **0.20** | **1.55 x 10^-3^** |
| **Morph (SC)** | **2.62** | **0.58** | **9.94 x 10^-6^** |
| Morph (YE) | -0.76 | 0.55 | 0.17 |
| **T_d_range_** | **2.50** | **0.27** | **< 2.0 x 10^-16^** |
| T_d_range_ x GH | -1.59 | 0.89 | 0.08 |
| **T_d_range_ x PS** | **-2.43** | **0.56** | **2.11 x 10^-5^** |
| T_d_range_ x SC | -0.33 | 0.84 | 0.11 |
| T_d_range_ x YE | -1.56 | 0.99 | 0.70 |

**Table S4.** Pairwise estimates of Weir and Cockerham weighted F_ST_ among populations used in acclimation treatments estimated from 1069 SNPs.

|  | *J. h. aikeni* | *J. h. dorsalis* | *J. p. palliatus* | *J. h. shufeldti* | *J. h. thurberi* |
| --- | --- | --- | --- | --- | --- |
| *J. h. aikeni* | 0 | 0.026072 | 0.051391 | 0.023624 | 0.026478 |
| *J. h. dorsalis* | 0.026072 | 0 | 0.028971 | 0.018739 | 0.01858 |
| *J. p. palliatus* | 0.051391 | 0.028971 | 0 | 0.051234 | 0.041593 |
| *J. h. shufeldti* | 0.023624 | 0.018739 | 0.051234 | 0 | 0.018673 |
| *J. h. thurberi* | 0.026478 | 0.01858 | 0.041593 | 0.018673 | 0 |

**Table S5.** Metadata for population genetic samples. Includes voucher catalogue number, species, subspecies, date collected, latitude and longitude of capture, NCBI Biosample, and barcode used in RAD library. Provided as .csv file.

**Table S6.** Acclimated individuals. Includes individual ID, population, latitude and longitude of capture, treatment, sex, adjustment period length, pre-acclimation mass, pre-acclimation M_sum_, time of pre-acclimation M_sum_ trial, mass post-acclimation, M_sum_ post-acclimation, time of post-acclimation M_sum_ trial, NCBI Biosample, and barcode used in RAD library. Provided as .csv file.

**Figure S1.** Approximate breeding ranges of *Junco* taxa in shaded polygons, derived from geo-referenced samples listed in Miller (1941). Dots denote origin of population genetic samples used in this study, with size of dot indicating number of specimens used for the corresponding locale (*n* = 1 to 4; detailed information in Table S5). Sample identification follows museum assignments..

**Figure S2.** Variance in M_sum_ among 106 wild-caught Oregon Juncos from Montana (*J. h. montanus*) decreased from $\sigma$ = 3.05 to $\sigma$ = 0.43 after a six-week adjustment period under common conditions in the lab (i.e. 18°C with 10 h:14 h light dark, *ad libitum* food and water). M_sum_ values collected within 24h of capture are included in the *in situ* dataset (Table S1). M_sum_ values after 6 wk adjustment period are published in Stager et al. 2020.
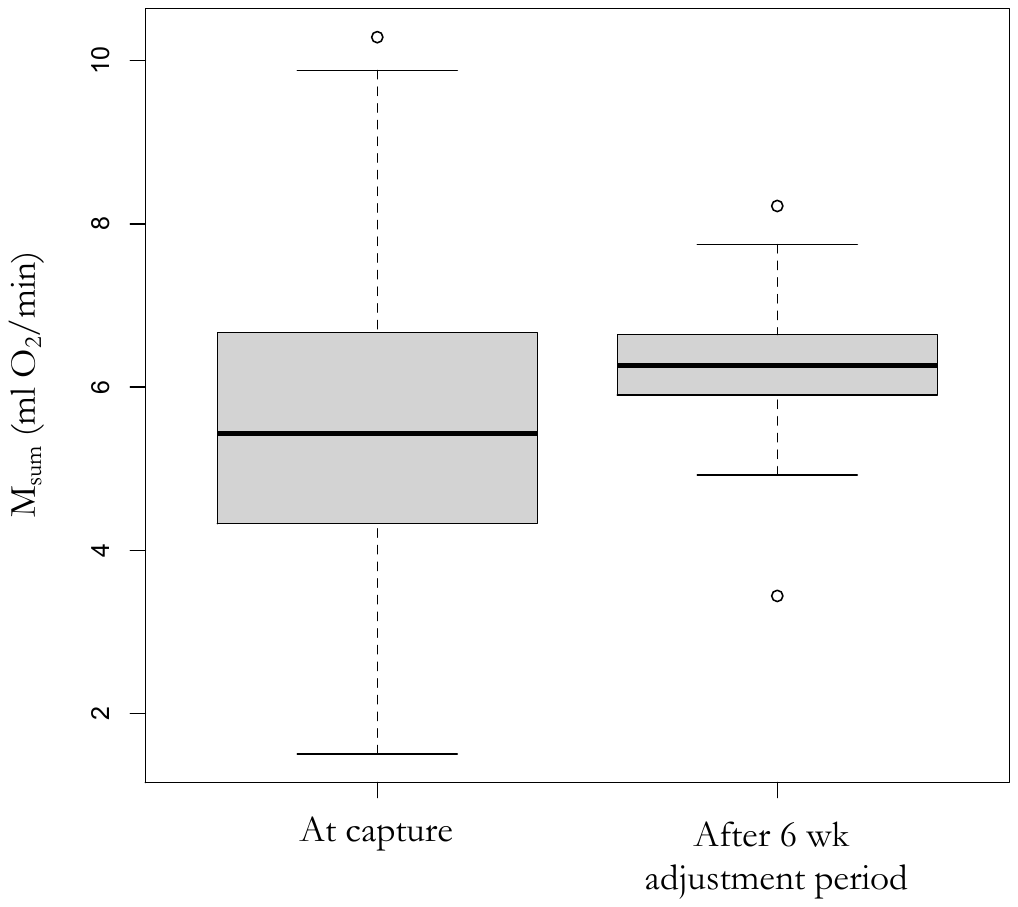
.

in a winter-tenacious songbird. *J Exp Biol* 223: jeb221853.
